# Antibody-based delivery of Interleukin-9 to neovascular structures: therapeutic evaluation in cancer and arthritis

**DOI:** 10.1101/2020.08.26.268292

**Authors:** Baptiste Gouyou, Tiziano Ongaro, Samuele Cazzamalli, Roberto De Luca, Anne Kerschenmeyer, Philippe Valet, Alessandra Villa, Dario Neri, Mattia Matasci

**Affiliations:** Philochem AG, Libernstrasse 3, 8112 Otelfingen, Switzerland; Department of Chemistry and Applied Biosciences, Swiss Federal Institute of Technology, Zurich, Switzerland; Institut des Maladies Métaboliques et Cardiovasculaires, INSERM U1048, Université de Toulouse, UPS, Toulouse, France

**Keywords:** Tumor targeting, Rheumatoid arthritis targeting, Fibronectin, Immunocytokines, Interleukin-9

## Abstract

Interleukin-9 (IL9) is a cytokine with multiple functions, including the ability to activate group 2 innate lymphoid cells (ILC2s), which has been postulated to be therapeutically active in mouse models of arthritis. Similarly, IL9 has been suggested to play an important role in tumor immunity. Here, we describe the cloning, expression and characterization of three fusion proteins based on murine IL9 and the F8 antibody, specific to the alternatively-spliced EDA domain of fibronectin. EDA is strongly expressed in cancer and in various arthritic conditions, while being undetectable in the majority of healthy organs. IL9-based fusion proteins with an irrelevant antibody specific to hen egg lysozyme served as negative control in our study. The fusion proteins were characterized by quantitative biodistribution analysis in tumor-bearing mice using radioiodinated protein preparations. The highest tumor uptake and best tumor:organ ratios were observed for a format, in which the IL9 moiety was flanked by two units of the F8 antibody in single-chain Fv format. Biological activity of IL9 was retained when the payload was fused to antibodies. However, the targeted delivery of IL9 to the disease site resulted in a modest anti-tumor activity in three different murine models of cancer (K1735M2, CT26 and F9), while no therapeutic benefit was observed in a collagen induced model of arthritis. Collectively, these results confirm the possibility to deliver IL9 to the site of disease but cast doubts about the alleged therapeutic activity of this cytokine in cancer and arthritis, which has been postulated in previous publications.

## INTRODUCTION

Interleukin-9 (IL9) is a multifunctional cytokine that has been originally identified as a T-cell and mast cells growth factor [1]. IL9 is secreted by mast cells and CD4+ T cells, including Th2, Th9, Th17, Treg, and NK cells [2] following stimulation by TGFβ and IL4 [3]. Its biological action is mediated by the heterodimeric receptor IL9-R comprising a specific IL-9 receptor α-chain and the common γ-chain cytokine receptor [4]. IL9 has been demonstrated to play an important role in a variety of physiological and pathological conditions including; pathogen immunity [5, 6], allergic inflammation [7, 8], and the immune regulation of cancer progression [9, 10].

The role of IL9 in tumor growth is controversial; whereas IL9 has been correlated to the progression of hematological malignancies both in mouse [11] and human [12], more recently IL9 dependent anti-cancer activity have been reported in a variety of solid tumor models including melanomas [13] and colon carcinomas [14, 15].

According to its postulated immunomodulatory activity, IL9 based therapies have been investigated in preclinical models of cancer and inflammation. In particular, Purwar and collaborators investigated the tumor immunity provided by IL9 in a preclinical therapeutic setting [13]. In the B16F10 melanoma models they demonstrated the ability of recombinant IL9 to impair tumor cell growth by a mechanism dependent on mast cells but not on T or B cells. The authors obtained similar results when recombinant IL9 was administered to mice bearing Lewis Lung carcinoma tumors suggesting a potential use of IL-9 based therapies for the treatment of diverse cancer types.

More recently, Rauber and collaborators, have identified IL9 as a key player in the resolution of rheumatoid arthritis in murine models of Antigen Induced Arthritis (AIA) and Serum Induced Arthritis (SIA) [16]. In their proposed mode of action, IL9 would promote ILC2 proliferation through an autocrine loop, finally resulting in the activation of Treg cells via a GITR/GITRL and ICOS/ICOSL dependent mechanism. The authors used a gene therapy approach based on hydrodynamic gene delivery to mediate recombinant expression of an IL9 cDNA construct in the treated mice. The *in vivo* levels of recombinant IL9 in arthritic mice were not measured, but a strong inhibition of disease progression, reduced tissue damage and bone loss was reported compared to control mice treated with a mock construct.

Collectively, the published work on the therapeutic use of IL9 prompted us to develop and evaluate new IL9-based immunocytokines and test them in preclinical models of cancer and arthritic diseases.

Various cytokine-based therapeutics may benefit from a targeted delivery to the site of disease, helping spare normal organs [17–19]. We have previously described an antibody-IL9 fusion protein and reported an unexpected impact of protein glycoform variation on tumor targeting performance [20]. Specifically, we used the F8 antibody, which recognizes the alternatively-spliced EDA domain of fibronectin, a marker of tumor angiogenesis and of tissue remodeling [21]. F8 reacts with murine and human EDA with identical affinity, since the target antigen features only three amino acid substitutions from mouse to man [21]. EDA is strongly expressed in the majority of solid tumors [22, 23], lymphomas [24] and in the bone marrow and chloroma lesions of acute leukemias [25] with a characteristic perivascular staining pattern, while the antigen is virtually undetectable in normal adult organs, exception made for the female reproductive system [26].

The tumor-targeting ability of the F8 antibody and of its derivatives has been established using radiolabeled protein preparations and quantitative biodistribution analysis [20, 21, 27–30]. Interestingly, pathological specimens from diseased tissues undergoing substantial remodeling (e.g., endometriosis, arthritis, psoriasis, atherosclerosis, vasculopathy and certain inflammatory bowel conditions) have been shown to be strongly positive for EDA(+)-fibronectin. The ability of F8 derivatives to preferentially localize at sites of chronic inflammation has been shown in mouse models [26, 27, 31–34] and in patients with rheumatoid arthritis [35].

In this article, we describe the design, expression and characterization of three recombinant formats of IL9-based fusion proteins, based on the EDA-targeting F8 antibody. IL9-fusions with the KSF antibody, specific for hen egg lysozyme, served as negative control with irrelevant specificity in the mouse. The fusion proteins were characterized in terms of their biochemical parameters, IL9 activity and tumor-targeting performance in biodistribution analysis after radioiodination. One of the formats, featuring the IL9 moiety flanked by two units of the F8 antibody in single-chain Fv (scFv) format, exhibited the best tumor targeting performance, with tumor:blood ratios > 10:1, twenty-four hours after intravenous administration. Whereas, a low increase of immune cells infiltration within tumors was observed, IL9 fusion proteins displayed only minimal therapeutic activity in three immunocompetent mouse models of cancer (K1735M2, CT26 and F9) at the maximal tolerated dose. Similarly, no retardation of disease progression was observed in the collagen-induced model of arthritis in mice. These results confirm that IL9 can be efficiently delivered to sites of disease with high expression of EDA *in vivo*, but cast doubts about the alleged translational therapeutic relevance of IL9 for the treatment of cancer and of chronic inflammatory conditions.

## MATERIALS AND METHODS

### Cloning of IL9 based fusion proteins

The genes encoding the antibody fusion proteins used in this study were generated by the assembly of multiple PCR fragments using as starting templates cDNA encoding for the anti EDA F8 antibody [21] or the irrelevant anti hen-lysozyme antib KSF antibody [36], either in scFv or diabody formats, and the murine IL9 (AA 19-144). The assembled constructs were inserted into the HindIII/NotI sites of the pcDNA3.1 (Invitrogen) mammalian expression vector. Primers used for the amplification and assembly of the different constructs are listened in the supplementary Table-1.

The pcDNA3.1-F8IL9F8 vector contains the murine IL9 sequence flanked at both the N-and C-termini by a 10-mer peptidic linker and the F8 antibody in scFv format. A leader sequence from murine IgG was used for protein secretion.

The pcDNA3.1-KSFIL9KSF contain a similar insert where the two F8 moieties were replaced by two copies of the KSF antibody in scFv format.

The pcDNA3.1-F8IL9 and pcDNA3.1-IL9F8, carry an insert consisting of the F8 antibody in diabody format fused either to the N-(pcDNA3.1-F8IL9) or C-(pcDNA3.1-IL9F8) terminus of IL9 via a 15-mer amino acidic linker. As well in these two vectors, a murine IgG leader sequence was used for protein secretion.

### Protein Expression, Purification and Characterization

The IL9 based fusion proteins (F8IL9F8, KSFIL9KSF, F8IL9 and IL9F8) were expressed by transient gene expression in CHO cells [37]. For transfection 4 × 10^6^ cells/mL cells were resuspended in ProCHO-4 Medium (Lonza) supplemented with 4 mM Ultraglutamine (Lonza). 0.625 μg of plasmid DNAs per million cells followed by 2.5 μg polyethyleneimine per million cells (Polysciences) were added to the cells and gently mixed. Transfected cultures were incubated in a shaking incubator at 31°C with 5% CO2 atmosphere shaking at 120 rpm for 6 days. The fusion proteins were purified by affinity chromatography using protein A affinity chromatography (Sino biological) according to the protocol provided by the supplier. Following elution proteins were finally dialyzed against PBS. Purified proteins were characterized for their size and homogeneity by SDS-PAGE and size exclusion chromatography, respectively. For SDS-PAGE analysis proteins were run under reducing and non-reducing conditions on 10 or 12% acrylamide gels (Invitrogen) and stained using Coomassie blue. Size-exclusion chromatography was performed on an ÄKTA FPLC system using a Superdex 200 increase 10/300GL column (GE Healthcare). Protein binding affinity was performed by Surface Plasmon Resonance using a BIAcore X100 instrument using a CM5 chip coated with recombinant fibronectin 11A12 domain. Samples were injected as serial-dilution, using a concentration range from 1mM to 125nM. The melting temperature of the different fusion proteins was determined using a StepOne real-Time PCR system (Applied Biosystems) in combination with the Protein Thermal Shift^™^ kit (Applied Biosystems).

### Immunofluorescence analysis of EDA expression

For biotinylation, 250 μg of fusion proteins were mixed to 50 μg of Biotin-NHS (Invitrogen) in a total volume of 1 mL. Reactions were gently agitated at room temperature for 1 hour prior to removal of unreacted Biotin-NHS by size exclusion chromatography using PD-10 columns (GE Healthcare). Elution was performed using PBS. Immunofluorescence staining was performed on three different frozen tissues: F9 teratocarcinoma tumors from 129/Sv mice, and healthy human basal and chorionic placenta. Frozen sections (8 μm) were fixed by ice-cold acetone and blocked with 20% fetal bovine serum in PBS. Detection of EDA positive fibronectin was performed using biotinylated fusion proteins followed by Streptavidin Alexa 488 (Invitrogen). In parallel the neovasculature marker CD31 was detected using either rat anti-mouse CD31 (BD Pharmingen) or mouse anti-human CD31 (eBiosciences) followed by Donkey Anti Rat-IgG-Alexa Fluor^TM^ 594 (Invitrogen), or Goat Anti Mouse-IgG-Alexa Fluor^TM^ 594 (Invitrogen), respectively.

### *In vitro* bioactivity assay

The bioactivity of mIL9 based fusion proteins was tested by a proliferation assay using MC/9 cells (ATCC). These mast cells derived from mouse liver were expanded in suspension in DMEM, supplemented with 10% FBS, 10% rat T-Stim, 2mM UltraGlutamine and 0.05mM–BetamercaptoEtOH. MC/9 cells were seeded at 40’000 cells per well in 96 wells plates in 200 μL of culturing medium (with 5% FBS and 2.5% rat T-Stim). Murine IL9 based fusion proteins and commercial murine IL9 (Peprotech) were added to the cells in serial dilution (from 28 nM until 0.1 pM). After 70 hours of incubation at 37°C with 5% CO2 atmosphere, 20 μL of Cell Titer Aqueous One Solution (Promega) was added to the wells and after incubation of 2 hours at 37°C the absorption at 492 nm was measured.

### Radiolabeled biodistribution and tumor therapies

*In vivo* experiments on tumor models were performed in agreement with the Swiss regulations and under a project license granted by the cantonal veterinary office (ZH027/15 for biodistribution and ZH004/18 for therapies). F9 teratocarcinoma cells (ATCC) were grown in DMEM (Life Technologies) supplemented with 10% FBS using 0.1% gelatin-coated tissue culture flasks. 129/Sv mice (8 weeks old, Charles River) were injected on the right flank with 13×10^6^ F9 teratocarcinomas. To assess tumor targeting in the F9 model, mice were grouped (n=3) when all tumors reached at least 200mm^3^. *In vivo* targeting was evaluated by biodistribution analysis as described previously [38]. 100 μg of each fusion protein were radioiodinated with ^125^I and Chloramine T hydrate and purified on a PD10 column (GE Healthcare), as described previously [38]. ~10 μg of radiolabeled proteins were injected into the lateral tail vein. Mice were sacrificed 24 h after injection. Organs were weighed and radioactivity was counted using a Packard Cobra gamma counter (Packard, Meriden, CT, USA). Values are given as percentage of the injected dose per gram of tissue (%ID/g ± standard error). K1735M2 melanomas were grown in DMEM (Life Technologies) supplemented with 10% FBS. C3H mice (8 weeks old, Janvier) were injected on the right flank with 5×10^6^ cells. Treatment started as soon as an average tumor size of >100 mm^3^ was reached, mice were randomized and grouped (n = 5 for IL9 based proteins, n = 4 for PBS). Mice were injected three times every second day with the test article intravenously (i.v.) in the lateral tail vein. 200 μg of F8IL9F8 or KSFIL9KSF were injected while PBS was injected at similar volume. CT26 colon carcinoma cells (ATCC) were grown in DMEM (Life Technologies) supplemented with 10% FBS. Balb/c mice (8 weeks old, Janvier) were injected on the right flank with 15×10^6^ cells. For therapies experiments with the CT26 cells, treatment started as soon as an average tumor size of >120mm^3^ was reached, mice were randomized and grouped (n = 5). Either 200 μg, or 100 μg of F8IL9F8 or equivalent volume of PBS were injected i.v. with the same schedule as the K1735M2 therapy. For F9 teratocarcinoma bearing mice, treatment started as soon as an average tumor size of >70 mm^3^ was reached, mice were randomized and grouped (n = 5). 200 μg F8IL9F8 or KSFIL9KSF were injected and the same volume of PBS was injected for the control group were injected i.v. with the same schedule as the K1735M2 therapy. For the three models, tumor volume was measured every day and the experiments were performed in a blind fashion. The therapy was discontinued by animal sacrifice when the weight loss exceeded −15% or tumor volumes was exceeding 1500 mm^3^. Data are expressed as the mean volume ± SEM. Differences between therapy groups were analyzed by two-way ANOVA multiple comparisons (Bonferroni corrected) using GraphPAD Prism (GraphPad Software, Inc.)

### Immunofluorescence analysis for *ex vivo* targeting and immune cells infiltrates studies

For FITC labeling, 1 mg of fusion protein was dialyzed in 0.1 M NaCO_3_ pH 9 then mixed with 25 μg of FITC (ACROS organics) in a total volume of 0.5 mL. Reactions were gently agitated at 4°C overnight prior to removal of unreacted FITC by size exclusion chromatography using PD-10 columns (GE Healthcare). Elution was performed using PBS. The *ex vivo* detection of the test articles was performed 24h after i.v. injection 160 μg of the FITC labeled proteins (F8IL9F8 and KSFIL9KSF) or PBS as control. Mice were sacrificed and tumors were OCT embedded. Frozen sections (8 μm) were fixed by ice-cold acetone and blocked with 20% fetal bovine serum in PBS. Detection of the test articles was performed using a rabbit anti-FITC IgG (Biorad) followed by anti-rabbit IgG Alexa Fluor^TM^ 488 (Invitrogen). The neovasculature marker CD31 was detected using a rat anti-mouse CD31 (BD Pharmingen) followed by Donkey Anti Rat-IgG-Alexa Fluor^TM^ 594 (Invitrogen). Sections were counter-stained with DAPI.

For the analysis of immune infiltrate into the tumors, mice received F8IL9F8 (n=2), KSFIL9KSF (n=1) or PBS (n=1) according to the same setting used in the therapy experiments (e.g. dose, schedule, tumor volume). 24h after the last injection, K1735M2 and CT26 bearing mice were sacrificed and tumor was OCT embedded. Frozen sections (8μm) were fixed by ice-cold acetone and blocked with 20% fetal bovine serum in PBS. Detection of the immune cells infiltrates were performed using rat anti-mCD8 (Biolegend), rat anti-mCD4 (Biolegend), rat anti-mFoxP3 (Invitrogen) or rat anti-mCD335 (NKp46) (Biolegend). Goat anti-rat IgG Alexa Fluor^TM^ 488 (Invitrogen) was used as detection antibody. Slides were counter stained with DAPI prior analysis. Sections were analyzed using an Axioskop2 mot plus microscope (Zeiss); pictures have been acquired with 10x magnification. For quantification of immune cells infiltrates 6 field of each section were analyzed, and the percentage of stained area was calculated by ImageJ.

### Rheumatoid arthritis therapy

For the collagen induced arthritis (CIA) experiments in mice; DBA/1J males (8-10 weeks old, Taconic Biosciences) were immunized by subcutaneous (s.c.) injection in the tail of an emulsion of bovine type II collagen in Completes Freund’s Adjuvant (CFA) (Hooke Laboratories). 18 days later, a booster injection of bovine collagen/IFA (Hooke Laboratories) was given to the mice by s.c. injection in the tail. Mice were scored every other day, the final CIA score represent the scoring sum of all 4 paws (0 = normal; 1 = one toe inflamed and swollen; 2 = more than one toe, but not entire paw inflamed and swollen or mild inflammation and swelling of entire paw; 3 = entire paw inflamed and swollen; and 4 = very inflamed and swollen paw). CIA scoring was performed blind. At the onset of the disease animals were randomized, before receiving their first injection. 150 μg, or 50 μg (n=15) of F8IL9F8 (n=15) or PBS (n=15) were injected in a final volume of 150 μL three times every second day i.v. in the lateral tail vein. Dexamethasone was daily injected intraperitoneally (i.p.) at 0.5 mg/kg.

## RESULTS

### All IL9 fusion proteins exhibit a good manufacturability profile

Different IL9 based fusion proteins were successfully expressed by transient gene expression (TGE) in Chinese hamster ovary (CHO) cells (Figure 1). The purified fusion proteins exhibited favorable biochemical properties as confirmed by SDS-PAGE and size exclusion chromatography (Figure 1 A-C). EDA-binding kinetic was analyzed by surface plasmon resonance (SPR). SPR analysis showed comparable nanomolar range apparent KD for all fusion proteins: 3.4 10^-9^ nM, 5.2 10^-9^ nM and 2.9 10^-9^ nM for F8IL9, IL9F8 and F8IL9F8 respectively (Figure 1 A-C). The bioactivity of the murine IL9 based fusion proteins has been assessed by their ability to stimulate the proliferation of MC/9 cells. In this assay, the three IL9 based fusion proteins showed similar activity with EC50 values of 38 pM (F8IL9F8), 21 pM (IL9F8) and 40 pM (F8IL9). The fusion of IL9 to the F8 antibody may slightly impair its function as a commercial mIL9 preparation showed a 10-fold higher biological activity (EC50 = 4pM). The comparative analysis of N-or C-terminal IL9 fusions showed that the arrangement of the different moieties did not affect its biological activity (Figure 1D). The single chain Fv based variant of the fusion protein (F8IL9F8) displayed a 3°C increased melting temperature compared to the two variants based on the diabody format (Figure 1 E), suggesting a better thermal stability. Indeed, when the F8IL9F8 variant was stored for up to 7 days at 4°C or 37°C or when tested over 3 cycles of freeze and thawing, no sign of aggregation, degradation or protein loss could be detected by size exclusion chromatography or SDS-PAGE analysis, (data not shown). Taken together these experiments suggest an overall better stability of the F8IL9F8 fusion protein.

**Figure 1:**
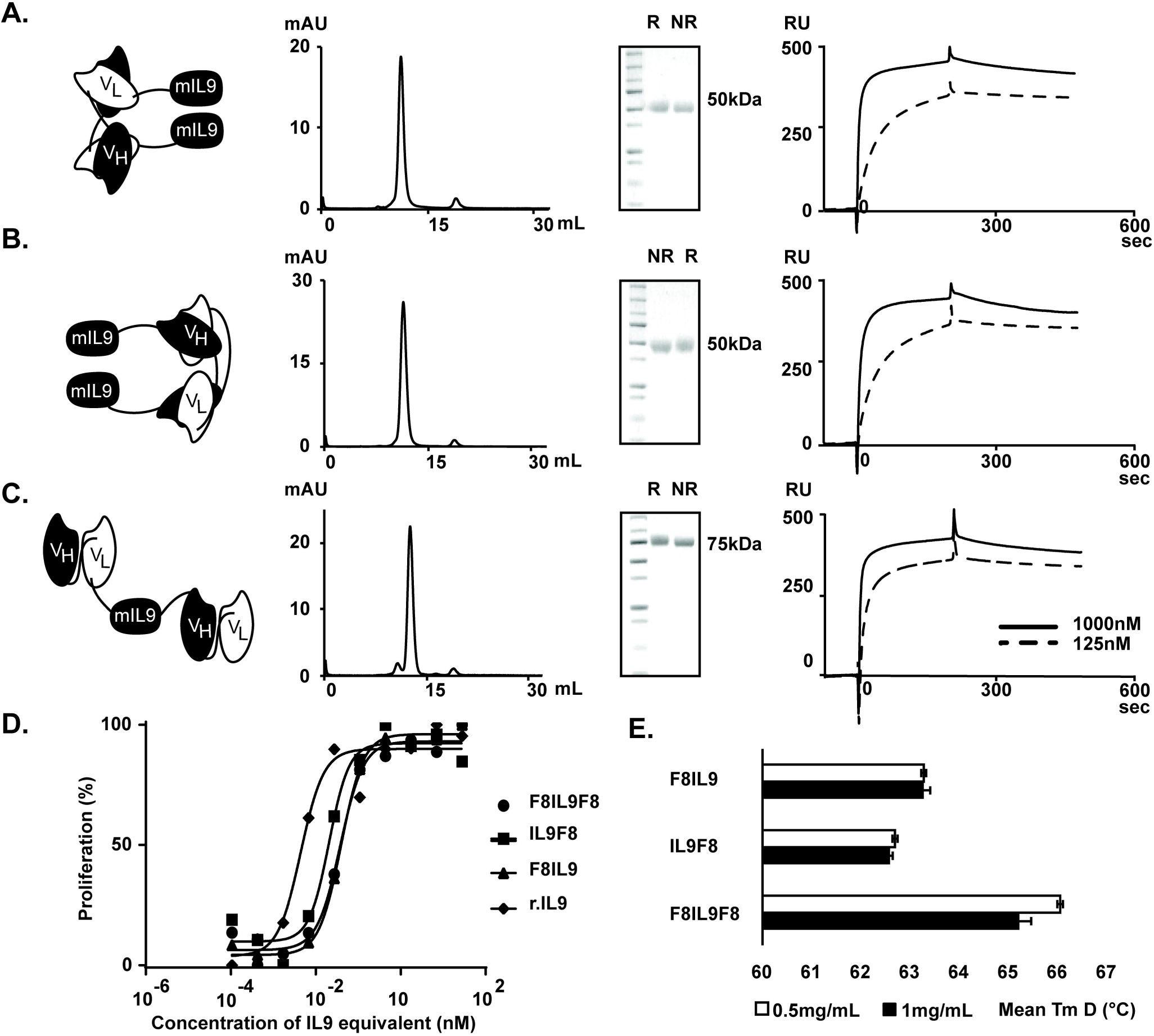
Cloning expression and characterization of IL9 based fusion proteins. IL9 fusion proteins have been expressed in CHO cells and characterized. From left to right: schematic representation of the IL9 based fusion protein, size exclusion chromatography profile, analytical SDS-PAGE analysis, SPR sensograms on EDA-coated sensor of (A) IL9F8, (B) F8IL9, (C) F8IL9F8. R: reducing conditions, NR: non-reducing conditions. (D) The proliferation of MC/9 has been assessed in presence of serial dilution of (□) IL9F8, (▲) F8IL9, (●) F8IL9F8 and (⍰) commercially available recombinant murine IL9 produced in bacteria. Data are presented as percentage of maximal proliferation (mean of three replicates). (E) Thermal shift assay was performed on IL9 based fusion protein. Data are expressed in °C + Std. Error, melting temperature (Tm) was assessed at two different protein concentrations.

### The format F8IL9F8 shows better *in vivo* targeting abilities

In order to test whether the IL9 fusion proteins retains the ability to bind EDA+ fibronectin on newly formed vascular structures, we performed immunofluorescence experiments on EDA positive tissue sections (i.e. human placenta and F9 tumors) [21]. The three fusion proteins selectively bound to neovasculature structures in both murine F9 teratocarcinoma and human placenta sections as confirmed by the co-localization with an anti-CD31 neovascular marker (Figure 2 E and F). Subsequently, the *in vivo* targeting performance of the three IL9 fusion proteins was tested by biodistribution analysis in tumor bearing mice. Purified fusion proteins were radiolabeled with Iodine 125 and 10 μg of each variant was injected into immunocompetent mice bearing F9 murine teratocarcinoma. The three variants were able to selectively localize to the tumor site at 24 hours post injection (Figure 2 A-D). The F8IL9F8 variant with two scFv antibody moieties was found to be superior when compared to the two other variants (IL9F8 and F8IL9, respectively) in terms of biodistribution profile, with an increased accumulation at the tumor site and at least a 2-fold increase in the tumor to organ ratios (Figure 2 A and B). For further *in vivo* investigation we decided to focus on F8IL9F8 due to its better targeting performance.

**Figure 2:**
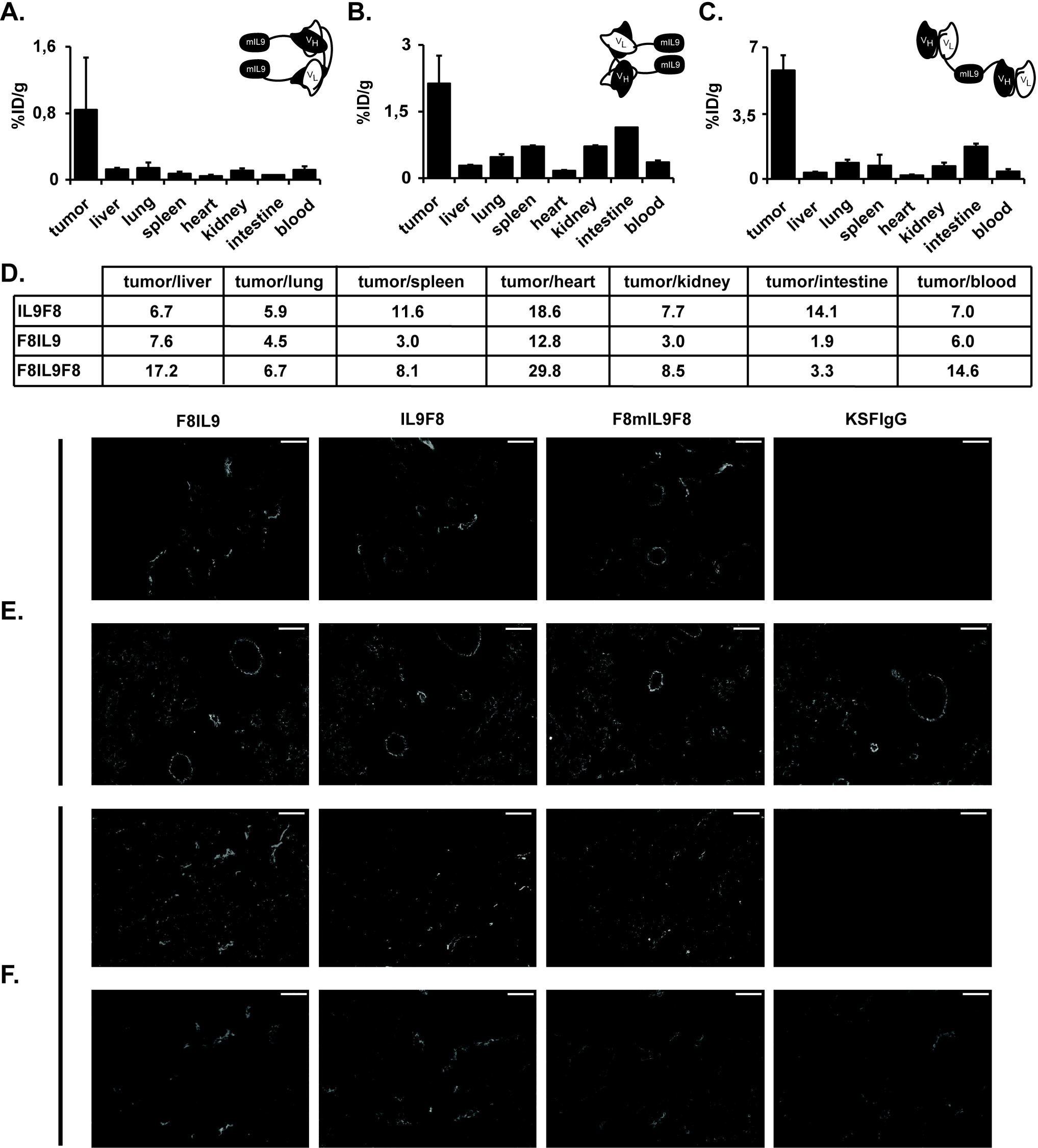
Tumor targeting performance of IL9-based immunocytokines. Quantitative biodistribution analysis of IL9 based fusion proteins. 10 μg of (A) IL9F8, (B) F8IL9 and (C) F8IL9F8 radiolabeled protein, have been injected into the lateral tail vein of F9 tumor bearing mice. Organs were collected, weighed, and the radioactivity measured 24 hours after injection. Results are expressed as percentage of injected dose per gram of tissue (% ID/g ± SEM), (n = 3 mice). (D) Table of tumor to organ ratios calculated from the average of the % ID/g. (E and F) Microscopic fluorescence analysis of (top panels) biotinylated IL9 based fusions proteins and (lower panels) anti CD31 antibody. Staining has been done against (E) healthy human placenta and (F) F9 Tumors grown in 129Sv mice, 10x magnification and scale bar represent 50μm. The irrelevant KSF antibody in IgG format, was used as negative control.

### IL9 fusion proteins display a therapeutic effect depending on cancer type tested

The therapeutic efficacy of F8IL9F8 was tested in three different cancer models: K1735M2 melanoma, CT26 colon carcinoma, and F9 teratocarcinoma.

Since previous studies showed the ability of IL9 or IL9-expressing Th9 cells to inhibit melanoma growth and reduce the number of melanoma-derived lung metastases in mouse [13, 39], F8IL9F8 was initially tested in the EDA-positive K1735M2 melanoma model [29]. Compared to the saline group, F8IL9F8 treatment exhibited minor tumor growth retardation which was statistically significant at the end point of the therapy (Figure 3A). Both F8IL9F8 and KSFIL9KSF treatments were well tolerated by mice, which displayed minimal and reversible weight loss. However, no substantial differences in tumor growth retardation and body weight change was observed between the treatments with targeted F8IL9F8 and the untargeted variant KSFIL9KSF (based on the irrelevant anti-hen egg lysozyme antibody KSF) (Figure 3A). The CT26 model, recently described as an immunologically “hot” tumor [40], was investigated due to earlier reports demonstrating *in vivo* growth impairment of CT26 tumors ectopically expressing IL9 [14, 15]. In order to investigate dose dependence effect of IL9 therapy, mice received three injections of F8IL9F8 at different doses, i.e. 100 μg or 200 μg. Both treatment regimens resulted in a similar tumor growth retardation when compared to the saline group, and no apparent signs of toxicity were reported (Figure 3B). However similar to K1735M2 melanoma also in the CT26 model no tumor eradication and mouse cure could be achieved with F8IL9F8. No therapeutic benefit of F9IL9F8 was observed in comparison to the saline group or mice treated with KSFIL9KSF in the F9 teratocarcinoma model. Interestingly, in this particular tumor model treatment with F8IL9F8 and KSFIL9KSF resulted in the highest body weight loss (Figure 3C).

**Figure 3:**
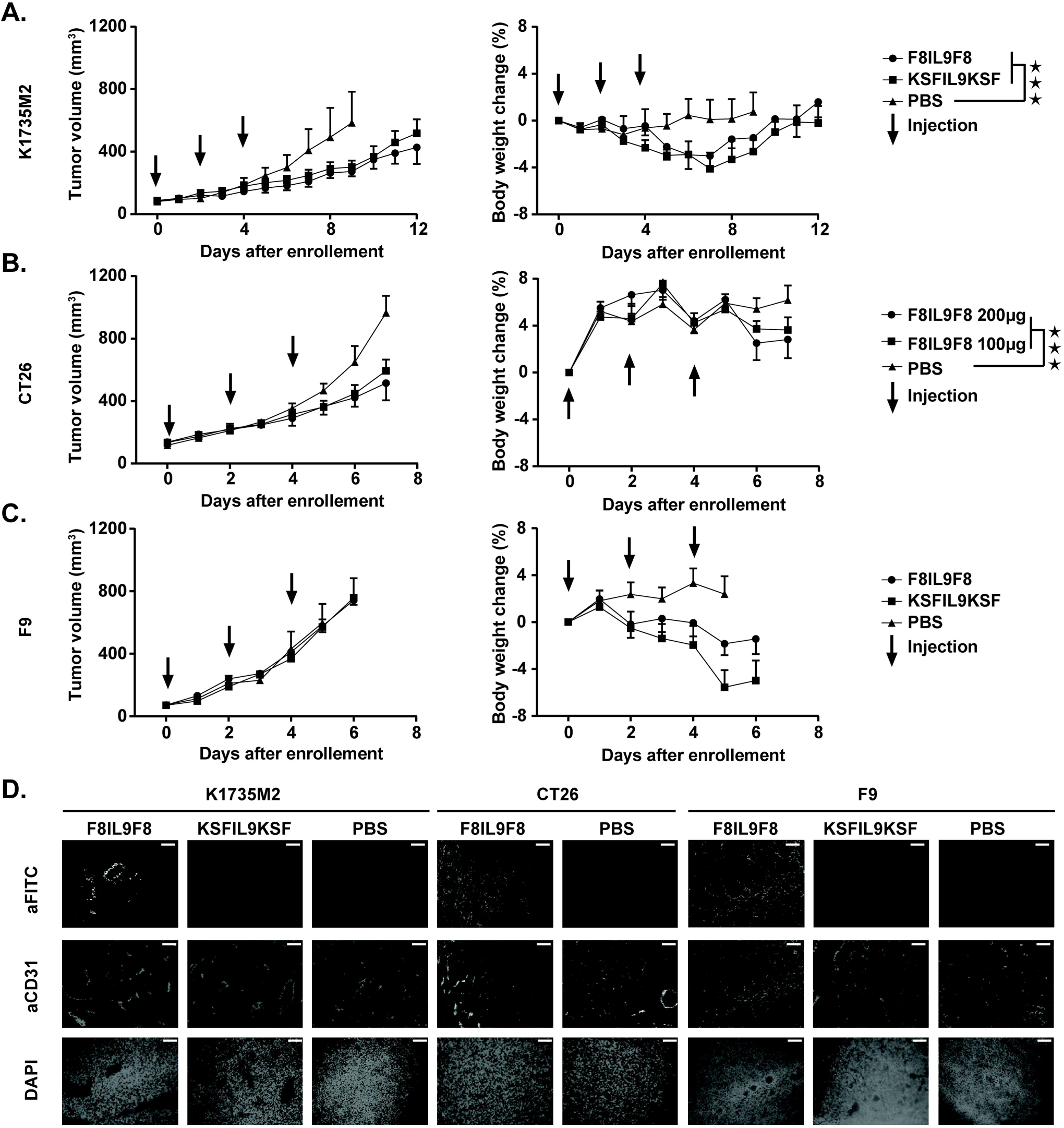
Therapeutic performance of IL9 based immunocytokines in C3H bearing K1735M2 melanoma, in Balb/c bearing CT26 colon carcinoma and in Sv129 bearing F9 teratocarcinoma. Comparison of F8IL9F8, KSFIL9KSF injected at 200 μg, and PBS, treatment started when K1735M2 reached an average volume of 100 mm^3^ borne in C3H (n = 5 mice for F8IL9F8 and KSFIL9KSF and n=4 for PBS) (A) Comparison of F8IL9F8 injected at 200 μg, at 100 μg and PBS, treatment started when CT26 tumors reached a volume of 120 mm^3^ borne in Balb/c mice (n = 5 mice per group) (B) and comparison of F8IL9F8, KSFIL9KSF injected at 200 μg, and PBS, treatment started when F9 tumors reached an average volume of 72 mm^3^ borne in Sv129 (n = 5 mice per group) (C). For all therapy experiments, mice were injected three times every second day i.v. in the lateral tail vein. Data represent mean tumor volume ± SEM (A-C left panel), body weight change ± SEM (A-C right panel). Statistical analysis was performed on tumor volumes using a 2-way Anova with a Bonferonni post-test ⍰: p-value < 0.05, ⍰ ⍰: p-value < 0.01, ⍰ ⍰ ⍰: p-value < 0.001, ns: not significant. Microscopic immunofluorescence analysis of the targeting ability of F8IL9F8, compared to KSFIL9KSF and PBS on the three tested cancer models, 10x magnification and scale bar represent 50 μm (D).

To confirm the *in vivo* targeting ability of F8IL9F8 in these therapies setting, *ex vivo* immunofluorescence studies were performed. In parallel to the therapy studies, additional K1735M2, CT26 and F9 tumor bearing mice were injected with FITC labeled fusion protein variants, sacrificed 24 hours and tumors were analyzed for the localization of the injected fusion proteins within the tumor mass. A preferential accumulation of F8IL9F8 around the neovasculature structures was observed for the three tumor models tested, by contrast no signal was observed for KSFIL9KSF or PBS injected mice, confirming the efficient and selective tumor targeting of the F8IL9F8 protein (Figure 3D).

### IL9 treatment is modifying the immune phenotype of tumors

The two models, responsive to IL9 therapy (i.e. K1735M2 and CT26) were analyzed for immune cells infiltrates at the end of therapeutic treatment. Multiple representative pictures for each treatment group were analyzed by ImageJ to quantify the percentage of positive area following staining for CD4+ T, CD8+ T, Treg and NK cells. A general increase in immune cells could be detected in both K1735M2 and CT26 tumors treated with IL9-fusion at 200 μg per injection (Figure 4), however to a degree considerably lower when compared to our previous reports with other targeted immunocytokines and different tumor models [30, 41–46]. Interestingly, in K1735M2 melanomas a comparable increment in CD4+T, and CD8+T cells was observed treated with either F8IL9F8 or KSFIL9KSF, whereas a slight increment in NK cells was observed only for F8IL9F8. Furthermore, no relevant changes in the infiltration of Treg cells could be observed between the 3 tested articles (Figure 4 A-C). Similar results were obtained for the CT26 colon carcinoma model, with a moderate increase of infiltrated CD4, CD8 and NK cells (Figure 4 B).

**Figure 4:**
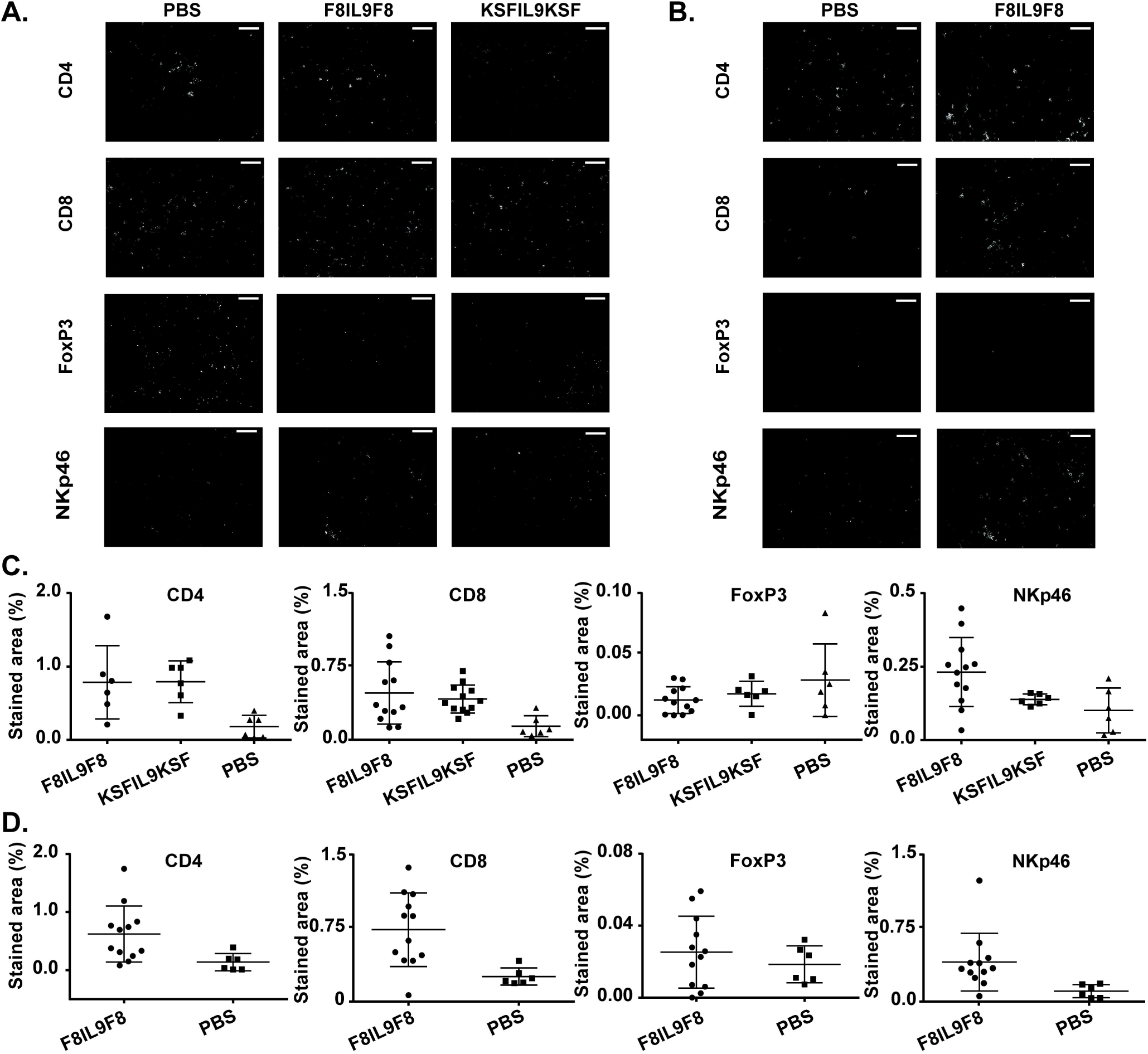
Immunofluorescence analysis on tumor-infiltrating immune cells in K1735M2 melanoma and CT26 colon carcinoma. Immunocompetent mice bearing K1735M2 (A and C) and CT26 (B and D), were injected three times every second day i.v. with 200 μg F8IL9F8, or 200 μg KSFIL9KSF or PBS. Mice were sacrificed 24h after the last injection and tumor sections were stained for CD4^+^ cells (CD4), CD8^+^ cells (CD8), Tregs (FoxP3) and NK cells (NKp46) (n = 2 mice for F8IL9F8 n=1 for KSFIL9KSF and PBS). Microscopic analysis of a representative section for each immunofluorescence, 10x magnification and scale bar represent 50 μm (A and B). Quantification of percentage of stained area using Image J. For each stained sections six representative areas were quantified and plotted as single dots, bars correspond to mean values with standard deviation (C and D). Full set of pictures used for quantification of area are represented in Figure S1.

### Delivery of IL-9 at the site of the disease did not provide a therapeutic benefit as an anti-inflammatory agent in collagen induced arthritis mouse model

The therapeutic potential of F8IL9F8 for the treatment of rheumatoid arthritis (RA) has been assessed in the CIA murine model. The disease was induced by two injections of a Bovine collagen preparation at distinct time points, which led to a rapid development of the disease. Mice were scored every other day using the standard scoring system. For each paw mice received 1 score for one toe inflamed and swollen, 2 for more than one toe, but not entire paw inflamed and swollen or mild inflammation and swollen paw, 3 for the entire paw inflamed and swollen; and 4 for very inflamed and swollen paw. At the onset of the disease, F8IL9F8 was injected either at high (150 μg/dose) or low (50 μg/dose) dose, and dexamethasone or PBS treatments were used as positive and negative controls, respectively. Whereas the positive control dexamethasone effectively reduced the diseases score already after 3 days from beginning of the treatment, the disease progression in the animal treated either with 50 or 150 μg/dose F8IL9F8 was comparable to the one of the mice in the saline group (Figure 5A). In agreement with the arthritic score increase, mice treated with F8IL9F8 or PBS showed a more prominent, yet reversible, body weight loss when compared to the positive control group treated with dexamethasone (Figure 5B).

**Figure 5:**
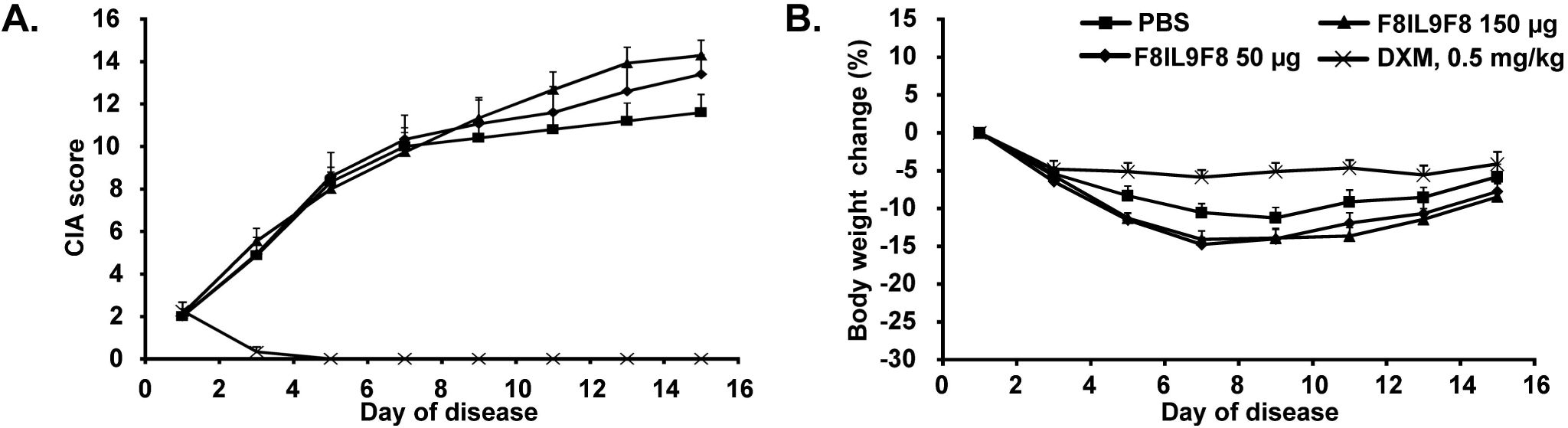
Therapeutic performance of F8IL9F8 in collagen induced arthritis (CIA) DBA/1 mice. Comparison of F8IL9F8, Vehicle (PBS) and dexamethasone treatment in collagen induced arthritis DBA/1 mice. At the onset of the disease (day 1) arthritic mice were injected three times every second day i.v. in the lateral tail vein with F8IL9F8 (150μg or 50μg) or Vehicle (PBS). dexamethasone was daily administrated (0.5mg/kg) i.p. from the onset of the disease. (n = 15 mice per group). Data represent from the onset of the disease mean CIA score ± SEM (A), body weight change from day 1 in percentage + SEM (B).

## DISCUSSION

We have generated and characterized three novel formats for antibody fusions with murine IL9. All three immunocytokine variants based on the F8 antibody, specific to the alternatively spliced EDA domain of fibronectin, could be expressed and purified to homogeneity. The products retained antigen-binding properties as well as *in vitro* IL9 biological activity. The F8IL9F8 product exhibited the best tumor targeting performance (6% ID/g in the tumor at 24h), with a tumor:blood ratio of 14.6. The percent injected dose per gram of tissue of IL9-fusions that could be delivered to tumors in mice was comparable to results obtained for other F8-fusion proteins with other cytokine payloads (e.g., IL2, IL4, IL6, IL10, IL12, TNF, interferon-alpha) [27, 30, 47–51]. However, unlike other F8 derivatives immunocytokines [27, 30, 47–51], F8IL9F8 did not display a potent anti-cancer activity *in vivo*. Similarly, F8IL9F8 was not active in a mouse model of arthritis.

The function of IL9 in tumor immunity is controversial [52–54]. It has been generally suggested that IL9 may inhibit the growth of solid tumors by activating the innate or adaptive immune response [55, 56], while as a lymphocyte growth factor it may promote progression of hematological malignancies [10, 12]. However, since IL9 has also been reported to promote proliferation of multiple solid tumor models (like pancreatic cancer, lung cancer and colitis associated cancer) [57–59], its anti-cancer activity may be context and tumor type specific [52–54].

In this study, we have used three syngeneic immunocompetent models of solid cancer known to express the EDA antigen (i.e. K1735M2 melanoma, CT26 colon carcinoma and F9 teratocarcinoma, respectively). When compared to saline treatment, a moderate anti-tumor effect could be observed *in vivo* only in the K1735M2 and CT26 models. However, targeting of F8IL9F8 did not result in improved therapeutic efficacy compared to the untargeted KSFIL9KSF at corresponding doses.

The antitumor function of IL9 on melanoma was initially postulated by Purwar and collaborators who showed that adoptive transfer of IL9-producing Th9 cells in mice bearing B16F10 melanoma tumors, was highly efficient in preventing tumor progression [13]. Accordingly, injection of recombinant IL9 resulted in impaired tumor growth, although to a lesser extent. More recently, Park and collaborators used B16F10 melanoma cells ectopically expressing soluble or membrane bound forms of IL9 to demonstrate the ability of IL9 to inhibit lung metastases [60]. A similar approach was used by Do Thi and collaborators. In this case, CT26 colon cancer cells engineered to ectopically express a membrane bound IL9, showed tumor growth retardation when compared to mock transfected CT26 cells [14, 15].

The exact mechanisms underlying IL9 mediated anti-tumor immunity are not fully understood. Whereas a prominent increase of CD4+ and CD8+ T and NK cells have been reported in tumor models ectopically expressing IL9 [14, 60], the work of Purwar suggest a key role played by mast cells since IL9 mediated tumor growth inhibition was abrogated in Kit W-sh mice (mast cells deficient mice) [13]. With our approach we expected that high IL9 concentrations at tumor site, resulting from the antibody-mediated targeted delivery of IL9, could contribute to a strong pro-inflammatory environment ideally favoring anti-cancer immunity and tumor eradication. Even though *in vivo* targeting of IL9 to the subendothelial matrix of the tested tumor models was successful, only moderate tumor growth retardation could be obtained. Similarly, only a very modest increase in tumor infiltrating CD4+ T, CD8+ T and NK cells could be observed in the two IL9 responsive models (i.e. K1735M2 and CT26). Similar findings have been reported by our groups: other F8 delivered payloads (e.g. IL17, IFNa and calreticulin) could increase immune cells infiltration without leading to noticeable anti-cancer efficacy in immunocompetent mice [36, 61, 62]. By contrast other immunocytokines (e.g. IL2, IL12, IL15, TNF) can promote strong neoplastic mass immune cells infiltration and strong anti-tumor efficacy [41, 45–47, 63, 64].

In a more recent study, Rauber and collaborators showed that IL9 could resolve chronic inflammation in two different models of arthritis (i.e. AIA in IL9 deficient mice and SIA models) [16]. They identified IL9-producting ILC2 cells as main mediators of the resolution of the chronic inflammation and proposed a mechanism by which IL9 would support ILC2 proliferation and survival by an autocrine loop. In our study, we did not observe a therapeutic benefit when F8IL9F8 was administered to arthritic mice. Both differences in the murine arthritis models and therapeutic approach used may account for the divergent results.

The CIA model was chosen since in this model we previously demonstrate strong EDA expression in the arthritic paws that were efficiently targeted by various F8 based immunocytokines [27, 31, 33]. Recombinant F8IL9F8 was administered after the onset of the disease in order to mimic as close as possible a potential clinical setting. By contrast, the studies of Rauber are based on non-standard therapy experiments, that would be difficult to be directly translated into clinical practice. IL9 was delivered by hydrodynamic gene transfer (HDGT) approach which do not allow control over the timing and concentration of overexpressed IL9. Furthermore, in their AIA study IL9 deficient mice were used, in which a chronic form of arthritis is developed, without signs of resolution. It should be noticed that whereas restoration of IL9 expression in the knockout mice could resolve chronic inflammation, no therapeutic benefit was observed in the wild type mice. Finally, in their serum induced arthritis model (SIA), IL9 delivery by HDGT was performed 5 days before disease induction, this would mimic a prophylactic rather than a therapeutic approach. The potential therapeutic benefit mediated by IL9 prior to the onset of the disease is in line with the recent finding that in the same SIA model, ILC2 adoptive transfer attenuated arthritis only if performed before SIA induction [65].

The therapeutic results obtained in the preclinical models of cancer and arthritis with F8IL9F8 were unexpected, particularly considering the previous results published by Purwar and Rauber [13, 16]. Although, it remains to be tested whether the anti-cancer and anti-arthritis activity of F8IL9F8 can be improved by combining it with other immunocytokines, or through a prophylactic approach, respectively. Currently, we do not believe that IL9 represent a suitable payload for the development of anti-tumor or anti-arthritis immunocytokines.

## ACKNOWLEDGEMENTS

We thank Catherine Pemberton-Ross and Lottie Howell for help with protein engineering and production.

Financial support by the ETH Zürich, the Swiss National Science Foundation (grant number 310030_182003/1), the European Research Council (ERC) under the European Union's Horizon 2020 research and innovation program (grant agreement 670603) is gratefully acknowledged.

