## Supporting information for "Antibody-based delivery of Interleukin-9 to neovascular structures: therapeutic evaluation in cancer and arthritis"

Table 1: Primers used for PCR amplification of IL9 based immunocytokines

| Payload | Fragment | Forward Primer | Backward Primer |
| --- | --- | --- | --- |
| F8IL9F8 | Fragment-1 | TCCTCCTGTTCCTCGTCGCTGTGGCTACAGGTGTGCACTCGGAGGTGCAGCTGTTGGAGTCTGGG | ACCTCCACCGCCAGAACCACTTCCGCCTGATTTGATTTCCACCTTGGTCCCTTG |
|  | Fragment-2 | TCAGGCGGAAGTGGTTCTGGCGGTGGAGGTCAGAGATGCAGCACCACATGGGG | CCCCCTGAACCACTGCCTCCAGATGGTCGGCTT |
|  | Fragment-3 | GCAGTGGTTCAGGGGGCGGTGGAGAGGTGCAGCTGT | TTTTCCTTTTGCGGCCGCTTATTTGATTTCCACCTTGGTCCC |
|  | Fragment-1/2 | CCCAAGCTTGTCGACCATGGGCTGGAGCCTGATCCTCCTGTTCCTCGTCGCTGTGGC | CCCCCTGAACCACTGCCTCCAGATGGTCGGCTT |
|  | Fragment-2/3 | CAGAGATGCAGCACCACATGGGGA | TTTTCCTTTTGCGGCCGCTTATTTGATTTCCACCTTGGTCCC |
| KSFIL9KSF | Fragment-1 | ATTAAAGCTTCCACCATGGGCTGGAGCCTGATCCTCCTGTTCCTCGTCGCTGTGGC | ACCGCCAGAGCCACCTCCGCCTGAACCGCCTCCACCACTCGAGACGGTGACCAGGG |
|  | Fragment-2 | AGGCGGTTCAGGCGGAGGTGGCTCTGGCGGTGGCGGATCGTCTGAGCTGACTCAGGA | CTCTGACCTCCACCGCCAGAACCACTTCCGCCTGAGCCTAGGACGGTCAGCTTGGT |
|  | Fragment-1/2 | ATTAAAGCTTCCACCATGGGCTGGAGCCTGATCCTCCTGTTCCTCGTCGCTGTGGC | CTCTGACCTCCACCGCCAGAACCACTTCCGCCTGAGCCTAGGACGGTCAGCTTGGT |
|  | Fragment-3 | ATGGGGAATTCGAGACACCAATTACCTTATTGAA | TAATGCGGCCGCTTATCATCCCAGCACTGTCAGCTT |

Table 1: (continued)

| F8IL9 | Fragment-1 | ATTAAAGCTTCCACCATGGGCTGGAGCCTGATCCTCCTGTTCCTCGTCGCTGTGGC | GCCAGAGCCACCTCCGCCTGAACCGCCTCCACCTTTGATTTCCACCTTGGTCCC |
| --- | --- | --- | --- |
|  | Fragment-2 | CAGGCGGAGGTGGCTCTGGCGGTGGCGGATCACAGGGGTGTCCAACCTTG | TAATGCGGCCGCTTATCATATCTTGCCTCTCATCCCTCT |
|  | Fragment-1/2 | ATTAAAGCTTCCACCATGGGCTGGAGCCTGATCCTCCTGTTCCTCGTCGCTGTGGC | TAATGCGGCCGCTTATCATATCTTGCCTCTCATCCCTCT |
| IL9F8 | Fragment-1 | TCCTCCTGTTCCTCGTCGCTGTGGCTACAGGTGTGCACTCGCAGAGATGCAGCACCACA | GCCAGAGCCACCTCCGCCTGAACCGCCTCCACCTGGTCGGCTTTTCTGCCTTTGCAT |
|  | Fragment-1' | ATTAAAGCTTCCACCATGGGCTGGAGCCTGATCCTCCTGTTCCTCGTCGCTGTGGC | GCCAGAGCCACCTCCGCCTGAACCGCCTCCACCTGGTCGGCTTTTCTGCCTTTGCAT |
|  | Fragment-2 | CAGGCGGAGGTGGCTCTGGCGGTGGCGGATCAGAGGTGCAGCTGTTGGAGTCT | TAATGCGGCCGCTTATCATTTGATTTCCACCTTGGTCCC |
|  | Fragment-1/2 | ATTAAAGCTTCCACCATGGGCTGGAGCCTGATCCTCCTGTTCCTCGTCGCTGTGGC | TAATGCGGCCGCTTATCATTTGATTTCCACCTTGGTCCC |

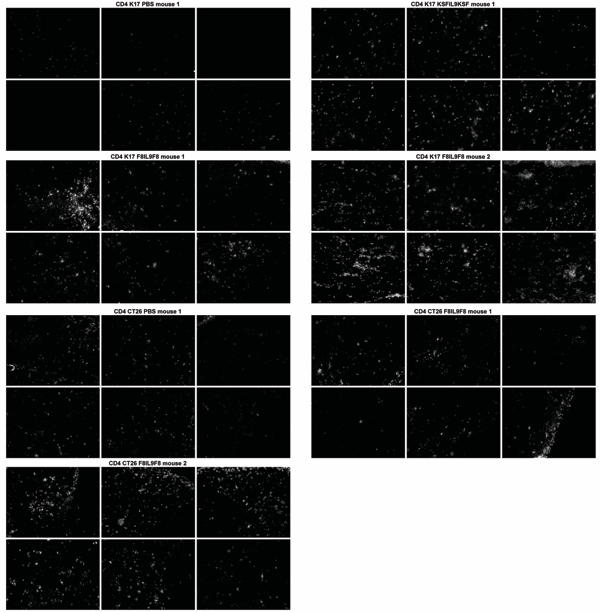

Figure S1: **Immunofluorescence analysis on tumor-infiltrating immune cells in K1735M2 melanoma and CT26 colon carcinoma.** Immunocompetent mice bearing K1735M2 and CT26, were injected three times every second day i.v. with 200 μg F8IL9F8, or 200 μg KSFIL9KSF or PBS. Mice were sacrificed 24h after the last injection and tumor sections were stained for CD4^+^ cells (CD4), CD8^+^ cells (CD8), Tregs (FoxP3) and NK cells (NKp46) (n = 2 mice for F8IL9F8, n=1 for KSFIL9KSF and PBS, 6 sections for each animal). Microscopic analysis of tumor infiltrating lymphocytes use for quantification via ImageJ.

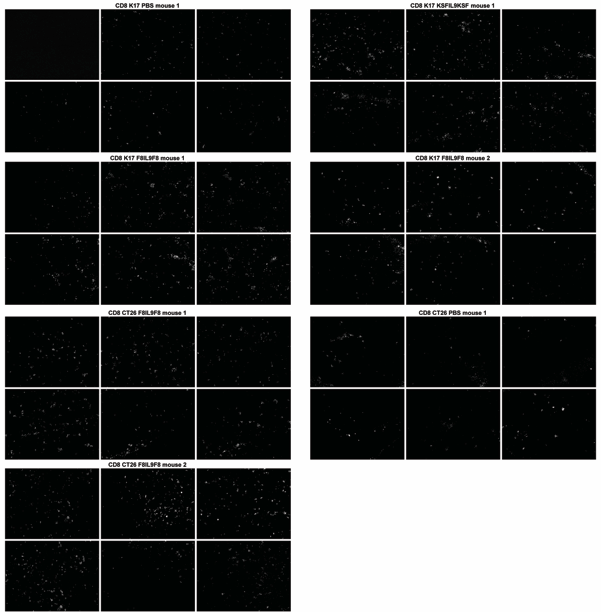
Figure S1: (continued)

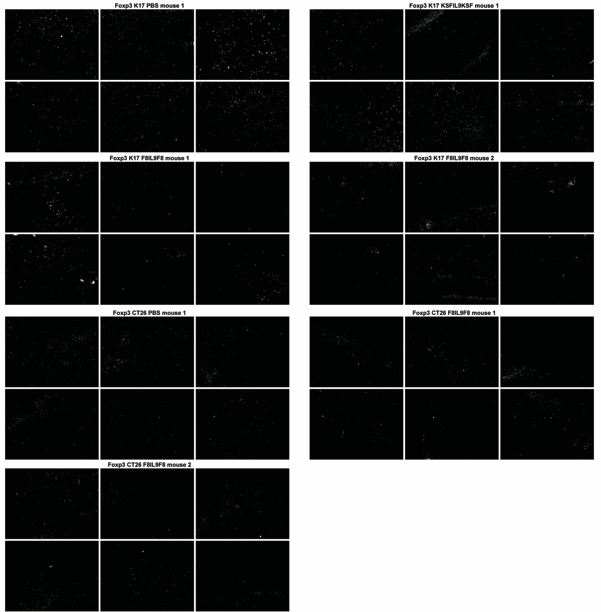

Figure S1: (continued)

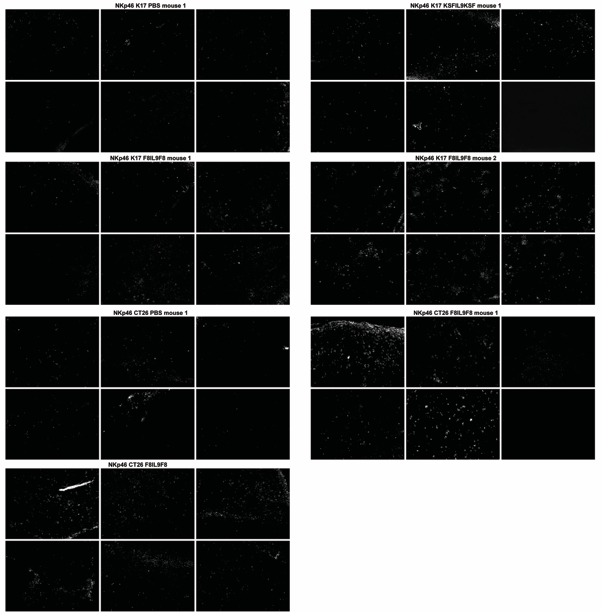

Figure S1: (continued)
